# Building a better frog trap: The benefits of mal-adaptive habitat choice for metapopulations with different life history strategies

**DOI:** 10.1101/004226

**Authors:** Rosemary Hartman, Noam Ross

## Abstract

By spatially distributing offspring among several habitat patches in varying environments, an organism can “hedge its bets” to protect against bad conditions in any single patch. This strategy can maintain populations even when some or even all locations are, on average, population sinks. However, species may not have evolved this bet-hedging mechanism, especially when sink environments are anthropogenic “traps” - locations where traditional habitat cues have been altered. Using a model based on the life history of the Cascades frog (*Rana cascadae*), we examine the conditions that maximize growth in an environment with ecological traps created by the introduction of predators. In a temporally stochastic environment, maximum growth rates occur when some juveniles disperse to the trap. We then examine how different life histories and predation regimes affect the ability of an organism gain an advantage by bet-hedging, and find that bet-hedging can be less useful when the ecological trap drives adult, rather than juvenile, mortality.

## Introduction

From deciding where to build a nest, deciding when to germinate, and deciding where to forage, organisms must attempt to use available cues to make habitat choices with important fitness consequences. Most models of habitat selection within a metapopulation assume that the organism has perfect information about habitat suitability (Delibes et al. 2001a). However, human impacts may change habitat quality in ways to which organisms have not adapted. Organisms continue to respond based on traditional cues, but these cues are no longer correlated to higher fitness. If enough individuals fall prey to these “ecological traps”, a human-impacted habitat patch may become a sink for the population (Dwernychuk and Boag 1972, Robertson and Hutton 2006, Schlaepfer et al. 2005). Modeling how maladaptive habitat selection affects metapopulation growth may reveal how human impacts affect sensitive species.

Many human actions can form ecological traps. Some well-known examples are resort lights on beaches which cause baby sea turtles to crawl the wrong direction upon hatching (Tuxbury and Slamon 2005), glass windows which elicit oviposition responses in mayflies, and floating garbage that resembles food to seabirds (Robertson et al 2013). Introduced species may cause inappropriate mating responses or habitat settlement cues (Schlaepfer et al. 2005). Introduced predators may be particularly problematic because native prey have not evolved to recognize them, and therefore fail to respond with anti-predator behaviors (Sih et al. 2010). If patches with predators are consistently superior in other habitat characteristics or resource availability (nesting sites, vegetation, water chemistry, prey food resources etc.) then ecological traps could cause the entire population to decline despite the presence of predator-free patches.

Sink habitats may also have value to organisms if they help maintain a population through risk-spreading or bet-hedging in a stochastically-varying environment (Holt 1997). (Throughout this paper we define “sink” as a habitat patch that could not persist in the absence of immigration.) Frequent changes in patch quality may mean that a patch that is a sink on average may convey higher fitness value at some times, facilitating metapopulation persistence (Holt 1997, Jansen and Yoshimura 1998). Some individuals settling in such patches may increase metapopulation growth and persistence more than all individuals settling in a single higher quality habitat, since temporary fluctuations in patch quality will affect a lower proportion of individuals.

Traditional models of source-sink dynamics assume passive dispersal, while most models that include habitat choice assume perfect information about habitat quality (reviewed by Piper 2011). When these assumptions are relaxed, population dynamics change. For example, Delibes et al. (2001b) used a deterministic model of active dispersal to examine how the equilibrium of a metapopulation changed as a function of the proportion of sink habitats, and showed a gradual increase in extinction probability where organisms had perfect information, but a sharp extinction threshold when the sink habitats were traps. However, environmental fluctuation may cause the bet-hedging predicted by Holt (1997), as even an ecological trap may have high fitness value during certain time periods. In these cases, a “trap” at the individual scale may not be a “trap” at the population scale.

Since traps can affect fitness at various life stages, life history may play an important role in whether mal-adaptive habitat choice at individual-scale translates to population-level declines. Differences in breeding site fidelity, natal site fidelity, parental care, and dispersal stage may change how a particular organism reacts to an ecological trap, and how the trap influences metapopulation dynamics. For example, organisms without parental care must choose habitat they believe to be suitable based purely on habitat cues rather than on prior breeding success (Blaustein 1999). The effect of this choice may depend on life history; some species may choose new breeding sites, while others remain or return year after year whether or not their young survive to adulthood (Refsnider et al. 2010).

To test how ecological traps affect metapopulations, we have constructed a theoretical model illustrating the dynamics of a two-patch metapopulation with mal-adaptive habitat choice. The model is based on the ecology of the Cascades frog (*Rana cascadae*), but potentially useful to aid thinking about a number of different organisms with ontogenetic differences in dispersal across a landscape of patchily distributed habitats. The basic model is best suited to describe other species with dispersing juvenile stages and sedentary adults, such as plants or marine invertebrates. Our extensions of the model explore how the model changes when a different life stage disperses, and how the model changes when predation affects the dispersing versus non-dispersing life stage.

We use this model to examine two questions. First, under what conditions can these ecological traps help maintain populations? Bet hedging theory would suggest that stochastic dynamics might make it beneficial for some individuals to disperse to lower-quality habitat patches if they change environmental state independently from the higher-quality patch (Philippi and Seger 1989). Secondly, we ask, how do differences in life history patterns affect the ability of organisms to capitalize on the bet-hedging strategy? Since longevity and iteroparity often buffer populations against temporal variability (Halpern et al. 2005), bet-hedging may be most important for species with short adult life spans.

In the following, we introduce a two-patch, two-stage, stochastic metapopulation model (“System Background and Model”). We analyze the effect of dispersal towards patches with predation on growth rate across different levels of predation and environmental variation (“Dispersal and growth”). We then examine the value of bet-hedging strategies under different life-history strategies, including different investment in survival of life stages, and the timing of dispersal, and the period of vulnerability to predation (“Life history scenarios”).

## System Background and Model

Cascades frogs live in high mountain lakes in the Cascades and Klamath Mountains in northwestern North America. They hatch and develop from tadpoles to frogs in a single season, then spend 2-3 years as juveniles while they disperse to other lakes. Adults choose a breeding lake at which they reproduce for the rest of their lives (Garwood 2009).

These lakes were historically fishless, but humans have introduced fish into some lakes, where they prey on frog larvae (Pope 2008). Previous studies have shown a negative relationship between fish presence and frog abundance (Welsh et al. 2006), but it is unclear whether this is due to predation alone or also to impacts of fish on adult frog habitat choice. Trout may cause an evolutionary trap for frogs choosing oviposiiton sites since they do not share recent evolutionary history and may not be able to recognize the presence of these novel predators (Sih et al 2010). Fish may decrease abundance of invertebrate predators (Pope et al 2009), which frogs have evolved to avoid (Hokit and Blaustein 1995). Therefore, it is possible that adult frogs will not recognize the presence of fish in a particular lake and oviposit there due to apparent lack of predators.

We represent this system with a two-patch metapopulation model with two life stages (juveniles and adults). Adults survive at rate *S*, reproduce at rate *f* and and juveniles recruit at patch-dependent rates *J*_*i*_. Juvenile recruitment rates in each patch are stochastic, with standard deviation *σ*_*Ji*_

One patch is an ecological trap with introduced fish (“predator patch”), and one is undisturbed (“predator-free patch”). Predators reduce juvenile recruitment by a factor of (1 *− p*). We consider predation pressure as an exogenous variable; though fish have a large effect on frog populations, frog larvae make up a very small percentage of fish diet so are unlikely to effect fish abundance (Joseph et al. 2011). Predation does not affect adult survival. For a single patch *i* with predation, the population projection matrix is:

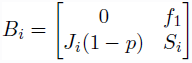

In the absence of predation, the (1 − *p*) term is removed.

We simplify the active process of habitat choice by individuals into a set fraction of individuals dispersing passively to each patch. All juveniles disperse, and a proportion of juveniles *d* settle in patch with predation. Juveniles are the only dispersing stage, and become adults once they settle in their destination patch. The dispersal matrix for the juvenile stage in a two-patch system is:

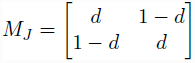

Combining these two matrices, the metapopulation projection matrix *A* for a system where demography occurs before dispersal is (adapted from Hunter and Caswell 2005):

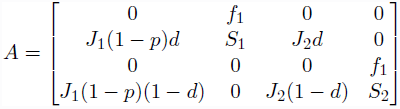

Populations in each patch are represented by the vector *n* = [*n*_*J*1_, *n*_*S*1_, *n*_*J*2_, *n*_*S*2_]. So the system’s evolution is represented as ***n***_*t*+1_ = ***An***_*t*_.

We drew life-history parameters from previous studies of *Rana cascadae* life history (Table 1) (Briggs and Storm 1970, Pope 2008).

**Table 1:** Vital rates for each life stage in a given environmental state. Juvenile recruitment was drawn at random from each of the four environmental states for each patch in each year. The “failure” state is one in which a bad water year causes very few juveniles to recruit into the population.

| Environmental state | Juvenile recruitment | Adult survival | Fecundity |
| --- | --- | --- | --- |
| good | 0.02 | 0.5 | 150 |
| average | 0.009 | 0.5 | 150 |
| bad | 0.002 | 0.5 | 150 |
| failure | 0.0001 | 0.5 | 150 |

All simulations were performed in R (R Core Team 2013), with some analyses using the popbio library (Stubben and Milligan, 2007). All code and simulated data are available at https://github.com/rosehartman/frog-trap.

## Dispersal and Growth

### Deterministic Growth

To see how changes in habitat selection affected population growth, we assumed density independent, deterministic growth and varied the dispersal to the patches with and without predation. To test the sensitivity of the metapopulation growth rate (log *λ*) to changes in predation and dispersal, we varied predation from 0% to 100% and varied proportion of juveniles dispersing to the predator patch from 0 to 1.

Under average-year conditions (table 1), with no inter-annual variation in survival (*σ_J__i_* = 0), any dispersal to the predator patch decreases the overall growth rate, an effect that is greater when predation was increased (Fig. 1). In a single-patch system, population growth rate of the predator patch would be positive at 0%, 20%, 40% and 60% predation, but negative (a true sink habitat) at 80% and 100% predation.

**Figure 1.**
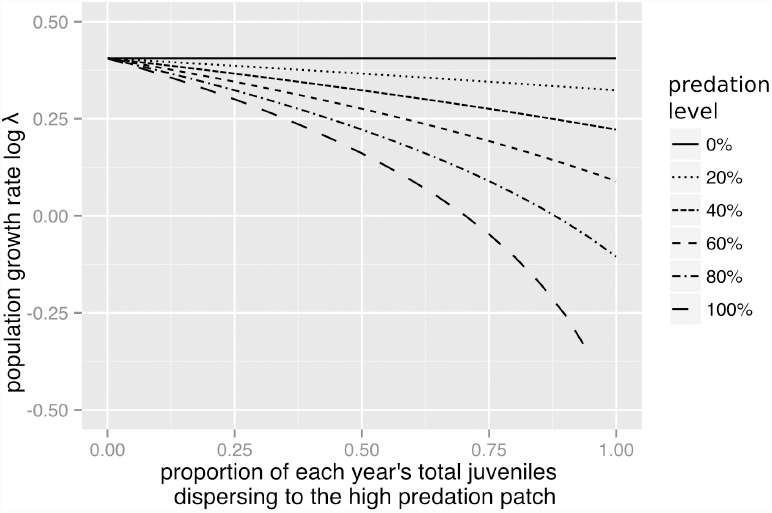
Growth rate of the two-patch metapopulation versus dispersal to the predator patch with deterministic survivorship.

### Stochastic Growth

We then introduced environmental stochasticity to the model by allowing each patch to be in one of four environmental states (Table 1). We have restricted variation in habitat quality to the juvenile life stage because juvenile recruitment is one of the most variable life history parameters in ranid frogs, and can be the major driver of fluctuations in population dynamics (Biek et al 2002). For frogs, local variation in habitat quality may be caused by abundance of vegetation, hydroperiod, predators, disease, large herbivores, or anthropogenic disturbance. The “failure” environmental state mimics a low water year or late freeze which cause very low juvenile recruitment.

Year types were chosen via a random draw based on equal probabilities of encountering each year type defined by different scenarios (Caswell 2001). We allowed the state in each patch to vary independently of the other patch with variation *σ*_*Ji*_, but juvenile recruitment in the predator patch was always penalized by the same percentage (p) no matter what the environmental state. We calculated stochastic metapopulation growth rates (log λ_*s*_) by averaging log λ for each year of one million year simulations. We varied predation percentages from 0% to 100% and the proportion of juveniles dispersing to the predator patch from 0 to 1. In a single-patch system, stochastic growth rate of the predator patch would be positive at 0%, 20%, and 40% predation, but negative at 60%, 80% and 100% predation.

Stochastic growth rate decreased with increasing predation. Increasing the dispersal to the predator patch increases the growth rate in a unimodel relationship (Fig. 2). The peak (marked with points on each curve) occurs near 50% of juveniles dispersing to the predator patch when predation is low, and moves towards lower values of *d* as the amount of predation increases. Even with 60% predation (sink habitat), some dispersal to the sink patch results in a greater log λ_*s*_ than total avoidance of the sink. However, if the patch is a very severe sink (80% or 100% predation), there is no longer a uni-modal relationship and log λ_*s*_ is maximized by total avoidance of the predator patch.

**Figure 2.**
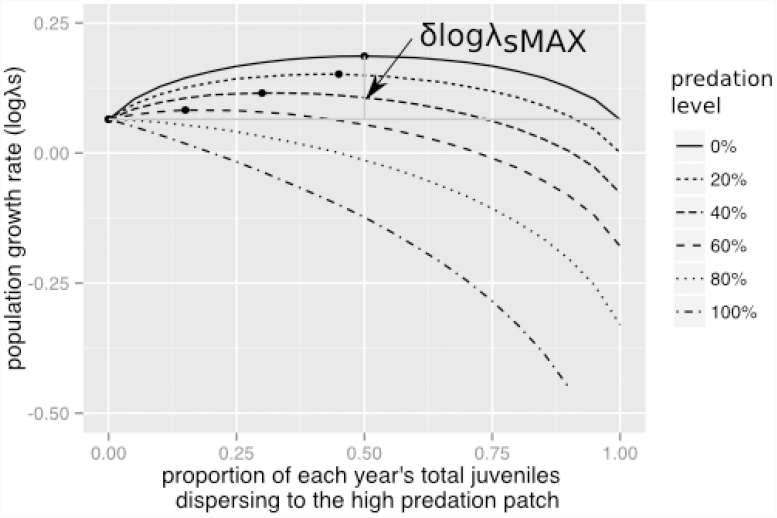
Growth rate of the two-patch metapopulation (log *λ*_*s*_) versus dispersal to the predator patch with stochastic survivorship. The maximum growth rate for each predation level (marked with points) occurs at lower dispersal toward the predator patch with increasing predation. The maximum increase in log *λ*_*s*_ (*δlogλ*_*sM A X*_, marked with vertical dotted line) decreases with increasing predation.

### Stochastic model analysis

To accompany our stochastic simulations, we analyzed the model, calculating stochastic growth rate using Doak et al’s (2005) modification of Tuljapurkar’s (1990) approximation:

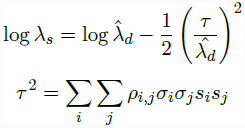

where 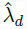 is the deterministic growth rate of the mean growth matrix, *i* and *j* are each of the parameters, *σ* are the standard deviations of stochastic parameters, *s* are the sensitivities of 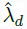 to each parameter, and *ρ*_*ij*_ the cross-parameter correlations. Tuljapurkar’s approximation assumes small noise, but has been shown to be robust (Fieberg and Ellner 2001).

For simplicity, in our analytic examinations we assume that *J*_1_ = *J*_2_ = *J*, and *σ*_*J*1_ = *σ*_*J*2_ = *σ_J_*. These changes do not have qualitative effects on our results.

We derived an expression for stochastic growth of the simplified model:

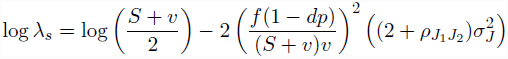

where

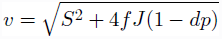

The term (*S*+ *v*)*/*2 is the deterministic growth rate under average conditions, or 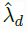. *S* can be interpreted as the contribution to growth of the current population, and *v* as the contribution of future recruits. 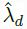 increases in *S*, *J*, and *f*, increasing fastest in *S*. The term *J*(1 *− dp*) in *v* represents the mean juvenile survival across both patches. *v*, and 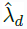, decreases with increasing *p* or *d*.

The second term in log *λ*_*s*_ is the change in the growth rate due to stochastic fluctuations, which is always negative. The fraction in the first parentheses generates the nonlinear relationship between *d* and log *λ*_*s*_. Both the *tau* component (*f*(1 *− dp*)*/v*) and the 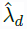 component (1*/*(*S* + *v*)) decrease with *d*, but at small values of *d*, the numerator decreases faster, leading to an increased growth rate, while at higher values of *d*, the denominator decreases faster, leading to a decrease in the growth rate. This can be interpreted as a decrease in the “stochastic penalty” for some amount of dispersion towards the predation patch, which is eventually overwhelmed by the lower average survival of juveniles in this patch. The stochastic penalty term also increases with larger stochastic fluctuations (*σ_J_*) and greater autocorrleation between fluctuations in the two patches (*ρJ*_1_*J*_2_).

To determine the level dispersal towards the predation patch, *d^∗^*that maximizes log *λ*_*max*_, we take the derivative of log *λ* with respect to *d* and set it to zero:

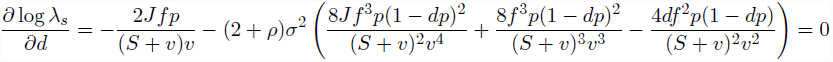

This expression is untractable for life history parameters other than *σ* and *ρ*. Our numerical results above indicate that *d^∗^* decreases with increasing *p*.

### Spatial autocorrelation

If patch environmental state is correlated, as in the case of nearby patches experiencing common climatic patterns, the value dispersal toward the predator patch changes. To see how this affected population growth rate, we varied the degree of spatial correlation between patches (*ρ*_*J*1*J*2_) from 0 to 1 and calculated the stochastic growth rate over one million years with a predation rate of 50% and varied proportion of juveniles dispersing to the predator patch from 0 to 1.

As the degree of spatial autocorrelation increases, the height of the dispersal-growth rate curve decreases, and the dispersal rate at which the curve peaks decreases (Fig. 3). This follows Equation 1 above, which shows log *λ* decreases with increasing *ρ*_*J*1*J*2_.

**Figure 3.**
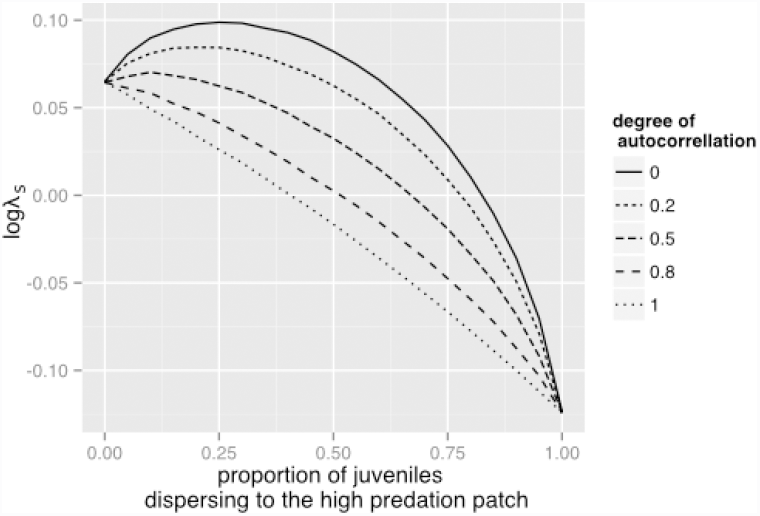
Dispersal-growth rate curve of metapopulation with varying degrees of spatial autocorrelation in year type. Predation was 50% in the predator patch. This is the average of 1000000 years.

## Life-history scenarios

The difference between the growth rate when juveniles always disperse to the predator-free patch (perfect habitat site selection), and the peak of the attractiveness-growth rate curve, can be thought of as the value of having some offspring make imperfect habitat choices. We will refer to this value as *δ* log λ_*s M A X*_.

### Relative survival of life stages

*δ* log *λ*_*s M A X*_ may change depending on the survival of the life stages that disperse. To examine the role of different life history strategies, we assume a trade-off between early life history stages and later life history stages. We multiplied the juvenile recruitment rate (independent of predation) for each environmental state by a factor of 0.1-2, while simultaneously changing the adult survival so that average life span (J/(1-S)) remains constant. Variation along this scale represents a trade-off in investment in early and late life stages. We simulated log *λs* as above, holding predation at 50%.

There is value in some dispersal toward a predator patch on the population level (the peak of the growth rate curve is greater than the growth rate at zero dispersal, see Fig 2). However, greater investment in adult survival decreases *δ* log *λ_s M A X_*, while greater investment in juvenile recruitment increased *δ*log *λ*_*s M A X*_(Fig 4, solid line).

### Timing of dispersal and predation

To expand our model beyond the life history of the Cascades frog, we tested sensitivity of log *λ*_*s*_ to changes in which life stage experienced predation and which life stage dispersed. We repeated the global elasticity analysis described above for an organism whose juveniles disperse but whose adults experience predation using metapopulation projection matrix A2:

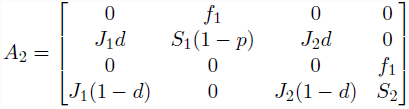

An organism whose adults disperse but whose juveniles experience predation is described by metapopulation projection matrix A3:

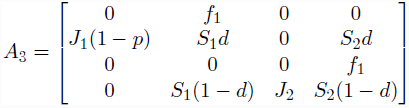

Lastly, an organism whose adults disperse and adults experience predation is described by metapopulation projection matrix A4:

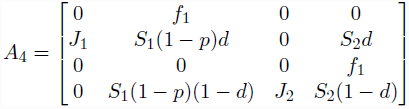

**Figure 4.**
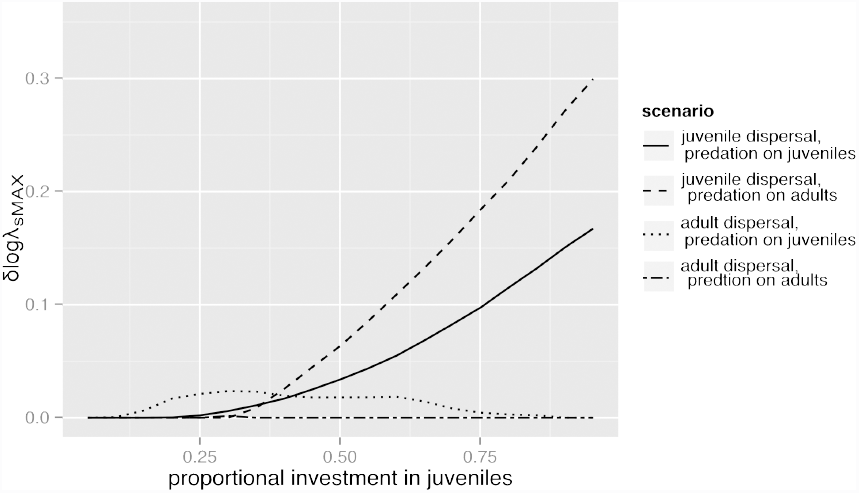
The gain from optimal dispersal towards the predation patch (*δ* log *λ*_*sM A X*_) across different life histories. In the baseline case (juvenile dispersal and predation on juveniles, solid lone), the gain increases with greater investment in juvenile survival. With predation on adults rather than juveniles (dotted line), there is a greater relative gain to be had by optimal dispersion towards the predation patch, again increasing with investment in juvenile survival. When adults disperse and but juveniles are preyed upon (dash-dot lines), the pattern is reversed, though log *λ*_*sMAX*_ is always low. When adults disperse and are preyed upon (dashed line), log *λ*_*sMAX*_= 0 in all cases.

Differences in life history change the magnitude of *δ* log *λ*_*sM A X*_. If adults experience predation instead of juveniles, but juveniles are still the dispersers, increases in adult survival still leads to lowering the fitness value of dispersal toward the predator patch (Fig. 4, dashed line). Allowing adults to disperse instead of juveniles causes a uni-modal relationship between adult survival and the fitness value of dispersing toward the predator patch (Fig 4, dotted line). However, for a given set of survival and fecundity parameters, the *δ* log *λ*_*sM A X*_ is lower when adults disperse than when juveniles disperse. When adults disperse every year and experience predation, there is no longer any value in dispersal toward a predator patch (*δ*log *λ*_*sMAX*_= 0, Fig. 4, dot-dash line).

## Discussion

### The value of ecological traps

While the presence of an ecological trap next to a high quality patch clearly decrease growth rates and equilibrium population densities in comparison to the presence of two high-quality patches, having some dispersal toward a low-quality patch is better for population growth and persistence than only having one patch. This can be caused by distribution of risk or bet-hedging in stochastically varying environments (Cohen 1966,Holt 1997). The degree to which dispersal toward an ecological trap increases population growth depends upon the difference in patch quality, the variance of the life history parameters, identity of the dispersing life stage, and the relative survivorship of the pre-dispersal and post-dispersal life stages.

The unimodal relationship between growth rate and attractiveness of the predator patch in the stochastic model is a response to uncertainty as to which patch will result in higher fitness. This can be thought of in the same terms as bet-hedging or risk-spreading (Philippi and Seger 1989), though the mechanism here is a proportion of juveniles mis-reading cues rather than adults intentionally ovipositing in multiple patches. Adults who have some offspring disperse to both patches will have higher average fitness than those whose offspring all choose the predator-free patch. While the predator-free patch has higher average juvenile recruitment, in any given year it may have lower juvenile recruitment than the predator patch.

There have been numerous other models showing how risk spreading may result in dispersal toward low-quality habitat. For example, a paper by Holt (1997) proposed that sinks are stable over evolutionary time because of high variance in survivorship at the source patch. While he also assumed organisms had perfect information on fitness in the two patches at the time they settled, the lack of knowledge of long-term dynamics makes preference for sink patches possible. Remes (2000) suggested that habitat choice, rather than dispersal limitation or despotic distributions, might actually drive the production of sinks when there was misinformation on the patch’s suitability. Hacco and Iwasha (1995) predicted that bet-hedging is most likely to evolve when there is poor information on environmental quality, such as the case of ecological traps that we explore here.

Previous models of ecological traps have not shown this uni-model relationship between growth rate and dispersal because they were either deterministic (e.g. Deleibes et. al. 2001), or because the model was parameterized in such a way that it was impossible for the sink patch to have occasionally higher survival than the source patch. In our model, when predation in the predator patch is 100% the predator-free patch will have higher juvenile recruitment in all year types (Fig. 2). In this scenario, the uni-model relationship disappears and there is no longer any benefit of having some dispersal toward the predator patch.

### Effect of spatial autocorrelation

Introducing a degree of spatial autocorrelation between the patches decreased the peak of the unimodal relationship between attractiveness of the trap and growth rate. It also significantly reduces the average growth rate for a given level of predation. This type of spatial synchrony makes the environment as a whole more predictable, because the predator patch always has lower fitness. Degree of spatial autocorrelation has previously been shown to determine amount of dispersal that optimizes population growth in source-sink metapopulation models (Schreiber 2010). More predictable environments make the strategy of high breeding site fidelity in this model less adaptive (Switzer 1993). If adults were able to switch patches based on previous breeding success, they could overcome the lower growth rates caused by the presence of the sink patch.

### Effect of stage-structured populations and life histories

The difference between the maximum metapopulation growth rate and the growth rate of the predator-free patch (log *λ*_*s*_*MAX*) represents the available gain from optimizing dispersal in a stochastically varying environment. Comparing patterns of log *λ_s_MAX* across life-history scenarios highlights how the effect of habitat choice varies.

Increasing investment in adults increased metapopulation survival more than an increase in juvenile investment (in most scenarios), however it decreased the value of having the ecological trap on the landscape (Fig. 4). Previous work has shown how adult longevity can buffer the effect of variability across time (Halpern et al. 2005), and may be a form of bet-hedging in itself (Saether and Bakke 2000). Environmental variability may also be the mechanism behind the evolution of iteroparity (Orzack and Tuljukupar 1989). When juveniles are the only dispersing life stage, increasing juvenile recruitment allowed more individuals to take advantage of a possible good year in the predator patch, thus increasing the value of the trap. Adults live for multiple years, so individuals who settle in the predator patch will have lower average fitness than those who settle in the predator-free patch. However, adults who have some offspring dispersing to both patches will have higher fitness than those whose offspring all go to one patch or the other. Therefore, an ecological trap may be more detrimental for a species with high adult survival than for a species with high juvenile survival.

When adults experience predation instead of juveniles, increasing adult survival decreases the value of dispersal toward a trap to an even greater degree, because the stochastic model is most sensitive to change in adult survival and increasing dispersal toward the patch with lower adult survival will decrease the average metapopulation adult survival. This type of system may be seen in harvested populations where only adults are harvested (Moffitt and Botsford 2009). No-take marine reserves or wildlife refuges provide “predator-free” patches, but dispersing juveniles may settle in areas where adults may be hunted.

In organisms with greater adult dispersal than juvenile dispersal, higher adult survival will increase the benefits of dispersal toward an ecological trap, since more frequent movements as adults allow more individuals to benefit from temporary good years in the predator patch. However, the benifit of dispersal toward a trap for these organisms is lower for one with juvenile dispersal because adults are not “stuck” in a trap for multiple generations. Multiple opportunities to switch patches makes initial dispersal decisions less important. The decrease in log *λ*_*s*_*MAX* at very high investment in adults show that when there is very low investment in juveniles the population may not be able to take advantage of the trap because it cannot grow fast enough to support itself. This life history strategy is seen in many large mobile vertebrates who have the opportunity to move their home ranges if a different patch appears higher in quality (reviewed by Piper 2011). Our analyses show ecological traps may be more beneficial for these species if their adults are long-lived with more opportunities to escape the trap or take advantage of temporarily better conditions.

However, when there is both adult dispersal and predation on adults, each individual adult will spend the same proportion of time in the trap habitat. Therefore, over a lifetime they will all have the same average fitness, negating the benefit of dispersal toward a trap. No uni-model curve is produced in this scenario, making it similar to previous models of ecological traps, such as Donovan and Thompson (2001) who found increasing dispersal rates of adults to low-quality habitat caused population extinction when percentage of low-quality habitat was greater than 30%. Previous models show presence of traps on the landscape, such as networks of marine reserves, drive selection toward greater site fidelity in adults (Meithe et al 2011), which may reflect evolution to escape from these types of traps.

Even when some dispersal toward a trap is beneficial, the genetic variation and/or phenotypic plasticity between individuals of the species will determine how many individuals disperse and whether the number who disperse toward the trap is optimal for maximizing the population growth rate (Clobert et al 2008). Because an dispersal to an ecological trap always decreases individual fitness over the long-term, the population benefit of dispersal to an ecological trap may not be evolutionarily stable. It will depend on whether the fitness benefit of having offspring disperse to both patches is greater than the fitness consequences of dispersing to the predator patch for the individual.

### Conservation implications

Ecological traps are often defined in terms of individual behavior; they are locations where cues are mismatched with habitat-selection behavior, often due to anthropogenic interference. However, such traps can nonetheless have value for growth or persistence at the metapopulation scale. Thus, in some cases the presence of ecological traps on the landscape may have conservation value. This would most likely occur where reduction in survival or fecundity in traps is less than complete, and where environmental variation in traps is not strongly correlated with variation in other habitats.

Importantly, the value of traps to the metapopulation depends on the life-history of the species. Species with high investment in juveniles, even if they are mal-adapted to new settlement cues, may gain benefit from some dispersal towards traps. Other species, especially those with longer-lived adults, may be unable to capitalize on this bet-hedging strategy at all.

Our model demonstrates limitations to the value of trap environments. The optimal dispersal toward the predator patch that maximizes metapopulation growth rate is always less than 50%. Therefore, if settlement cues in the traps are much greater than in other environments, so that most or all individuals fall into this trap, they will not increase population growth or persistence.

In those cases where traps do have potential value, it is possible that strategies such as removing settlement cues, erecting barriers or resettlement to habitats with environments with higher average survivorship will be less effective than strategies that improve the quality of trap habitat or reversing anthropogenic changes that created traps.

Traps may also be mediated through changes in the organisms themselves. They may escape by rapid evolution, phenotypic plasticity, or philopatry (Kokko and Sutherland. 2001, Meithe et al 2011), and knowing whether any of these can occur may be key to managing populations in a changing environment.

## Conclusions

Maladaptive habitat selection decrease does population growth more than presence of low-quality habitat patches on their own, but stochastic dynamics and non-linear growth curves may mean a slightly attractive sink is better than one that is totally avoided. Specifics of the organisms’ life history, dispersal rates, and spatial variation in factors affecting survival are key to determining the degree to which ecological traps are helpful or detrimental to population persistence.

